# Targeted Esterase induced Dye loading supports Calcium Imaging in Eukaryotic Cell-Free Systems

**DOI:** 10.1101/2020.03.05.978247

**Authors:** Priyavathi Dhandapani, Srujan Kumar Dondapati, Anne Zemella, Dennis Bräuer, Doreen Anja Wüstenhagen, Stefan Kubick

## Abstract

Calcium imaging is an important functional tool for addressing ion channels, transporters and pumps for drug screening in living cells. Depicted eukaryotic cell-free systems utilize microsomes, derived from endoplasmic reticulum to incorporate the synthesized membrane proteins. Absence or inadequate amount of carboxylesterase in the endoplasmic reticulum of eukaryotic cells, which is necessary to cleave the acetoxymethyl ester moiety of the chemical calcium indicators, advocates the hindrance to perform calcium imaging in microsomes. In this work, we try to overcome this drawback and adapt the cell-based calcium imaging principle to a cell-free protein synthesis platform. Carboxylesterase synthesized in a Spodoptera frugiperda Sf21 lysate translation system is established as a viable calcium imaging tool and hTRPV1 is used as a model channel protein to demonstrate the realization of this concept.

## INTRODUCTION

Calcium permeable membrane proteins such as ion channels, transporters and pumps, contribute to the majority of eukaryotic membrane proteins, serving as viable drug targets for several pathological diseases next to the large family of G-protein coupled receptors (1). Eukaryotic cell-free protein translation overcomes several disadvantages that could be met for overexpression of ion channels, transporters and pumps such as cell-toxicity, poor expression and deletion due to engineered protein domains (2). As eukaryotic cell-free translated membrane proteins are incorporated into microsomal membranes and removal of the plasma membrane during lysate preparation grants us with numerous advantages of working with plasma membrane channels in the microsomes. One big advantage is that increased signal to noise ratio in assays over cell-based expression. In cell-based protein expression, ion channels are evaluated using non-functional assays such as ligand binding assays and functional assays such as automated and manual electrophysiology as well as voltage and ion sensitive dye binding assays. Although many researchers have worked on ion channels synthesized using prokaryotic cell-free synthesis, little work on ion channel expression and functionality with eukaryotic systems have been done till date. Some of them are: planar bilayer measurements such as alpha7 nicotinic acetylcholine receptor (nAChR)(3), connexion gap junction channels (4) in a rabbit reticulocyte lysate translation system, (human voltage dependent anionic channel (hVDAC1) (5), K channel of streptomyces A (KcSA) (6), Bombyx mori olfactory receptor BmOR1 (7) in a Spodoptera frugiperda *Sf*21 lysate translation system, shaker potassium channels (8) in the wheat germ cell extract translation system; membrane topogenesis studies in microsomes such as Inward-rectifier potassium channel (Kir 2.1) and KcsA (9) in the rabbit reticulocyte system.

Although some current evidences have been reported for electrophysiological experiments, no evidence on functionality assays with ion and voltage sensitive dyes is currently available. The concept of calcium imaging of eukaryotic cell-free translated channels revolves around the previously reported idea of Targeted Esterase induced Dye loading (TED) of the endoplasmic reticulum (ER) in mammalian cells (10, 11). We use a mouse carboxylesterase 2 (*m*CES2) variant because, it is already well characterized for low affinity chemical fluorescent calcium dyes (10). We focus on studying calcium, being customarily a permeabilized ion through several families of ion channels and a second messenger in signal transduction in large number of cellular pathways. Intraluminal calcium is the prime regulator for ER function, which has also been studied concomitantly in microsomes by researchers in last decades. Using non-fluorescent methods such as radioactive labelling of calcium ions, extensive studies have been reported using rat brain microsomes (12, 13), hepatic microsomes (14) and rabbit skeletal microsomes (15). All the aforementioned radioactive Ca^2+^ uptake and release studies have been performed only for native channels and pumps present in the ER such as Sarcoplasmic reticulum Calcium ATPase (SERCA) and leak channels such as Ryanodine receptors (RyR1 and RyR2).

## RESULTS AND DISCUSSION

### Cell-free synthesis using Sf21 system

In order to establish a vesicle-based calcium imaging tool, *m*CES2 cell-free construct was used as depicted in Fig 1 c to incorporate the soluble protein *m*CES2 inside the lumen of microsomes during cell-free translation. A melittin signal sequence upstream of the gene was utilized as aforementioned (16). As the model protein *h*TRPV1 is a poly transmembrane channel, no signal sequence is necessary for active incorporation into microsomal membranes. A CrPV IRES site upstream of the gene for both *m*CES2 and *h*TRPV1 was used with a start codon as GCT, alanine instead of ATG, methionine for improved translation as reported before for other proteins (17). T7 promoter and T7 terminator was used for mRNA transcription. For cell-free synthesis of *m*CES2 and *h*TRPV1, we used the Continuous Exchange Cell-Free (CECF) mode of our *Sf*21 translation system (18,19,20). This mode of synthesis has mainly the amino acids and the energy components in the larger compartment and the eukaryotic translation mix in a smaller compartment separated by a dialysis membrane of 9 kDa cut off. Caspase inhibitor was used for improved yields and avoiding protein degradation during the incubation step of 24 h. Though the total translation mix contained 85 ng/μl of the *de novo* synthesized *m*CES2 protein, only 25 ng/μl was obtained in the vesicular fraction. For expression of both the proteins, *h*TRPV1 and *m*CES2 were correspondingly sequentially translated for 24 h each. As the microsomes obtained from the first translation were used in the second translation, it may occur that the first translated protein is not present in all the microsomes at the end of the second translation. To avoid this issue, we have used only the vesicular fraction obtained after second translation for all our experiments that require tandem protein translation. Simultaneous translation is not preferred as evaluating the amount of individual proteins synthesized is not feasible. *m*CES2 and *h*TRPV1 synthesized showed a molecular weight of 58 kDa and 98 kDa correspondingly under reduced conditions. As *m*CES2 and *h*TRPV1 contain di-sulphide bridges, prudently the translation is synthesized only under non-reducing conditions.

**Fig 1:**
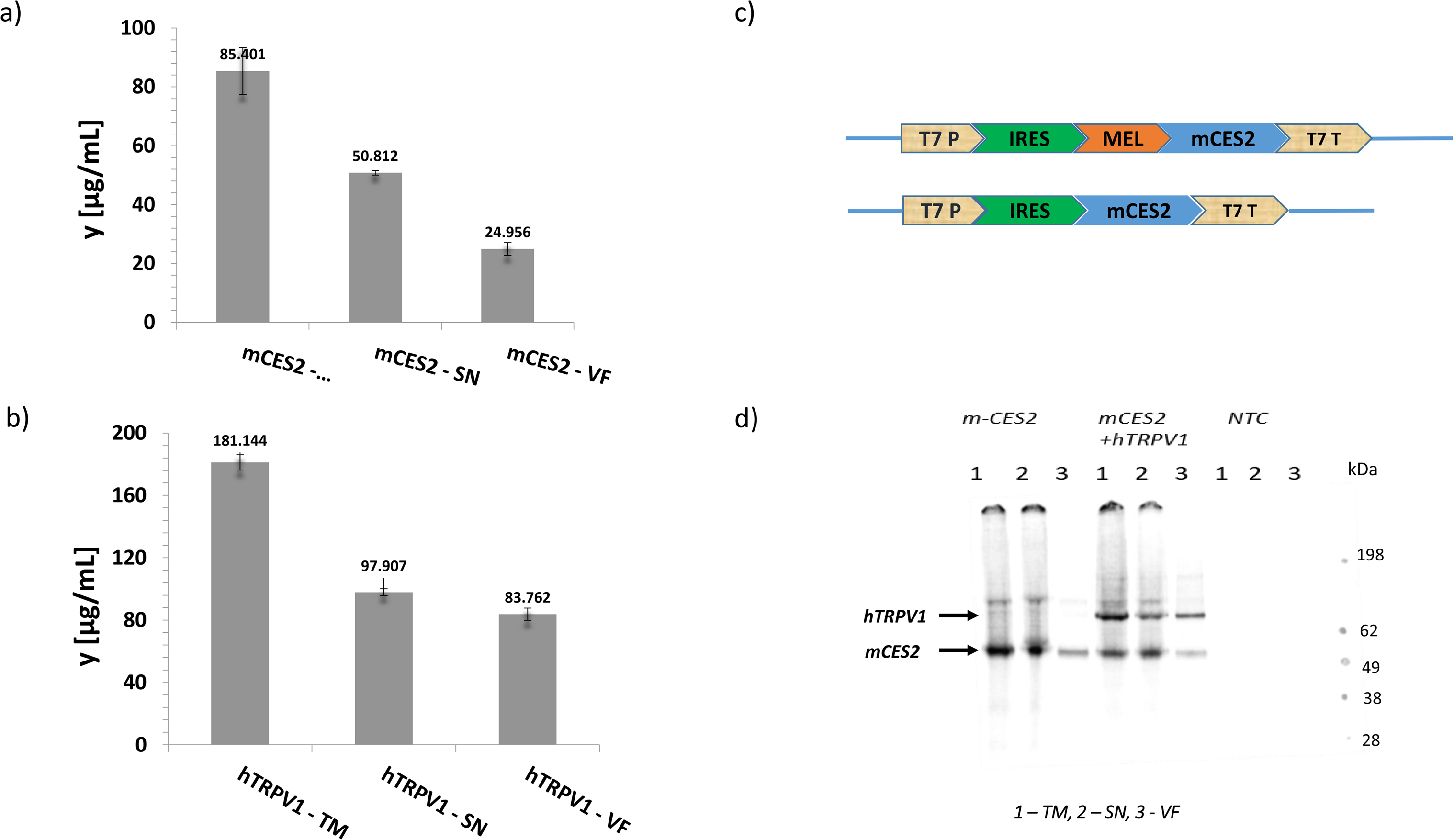
Cell-free protein synthesis of *h*TRPV1 and *m*CES2. a) Protein yields of *m*CES2 by scintillation counting via CECF reaction for 24 h, 30 °C in Sf21 system; b) Protein yields of hTRPV1 by scintillation counting via CECF reaction for 24 h, 30 °C in *Sf*21 system c) Template design of *m*CES2 and *h*TRPV1, MEL-melittin signal sequence, T7 P - T7 promoter, T7 T – T7 terminator, IRES- Cricket paralysis virus Internal Ribosome Entry Site, d) Autoradiogram of proteins run on SDS MES

### Validation of synthesized *m*CES2 in microsomal lumen

Carboxylesterases are present in the cytosol in large quantities, which also results in the lysate. Apparently, the cytosolic carboxylesterase is likely to be present outside the microsome and will not contribute for any applications like calcium imaging of microsomes. Prior to functional assessment, intensive washing of microsomes was performed to remove the cytosolic esterase. Functionality of the *m*CES2 synthesized inside the microsomes is assessed using the commonly used para-nitrophenyl acetate (PNPA) method (21, 22) (Fig 2). The PNPA method provides a faster way to assess the esterase activity. The esterase activity is preferably analysed for maximum 1 h in accordance with the time frame for calcium imaging applications. *m*CES2 microsomes showed higher esterase activity relative to NTC microsomes in a substrate dependent (Fig 2 b) plot, time dependent and dose dependent (Fig 2 d, supplementary data 1) measurements. The activity observed in the NTC microsomes is probably due to unspecific activity of hydroxylase or a mono-oxygenase class of cytochrome P450 enzymes present in the endoplasmic reticulum (23, 24).

**Fig 2:**
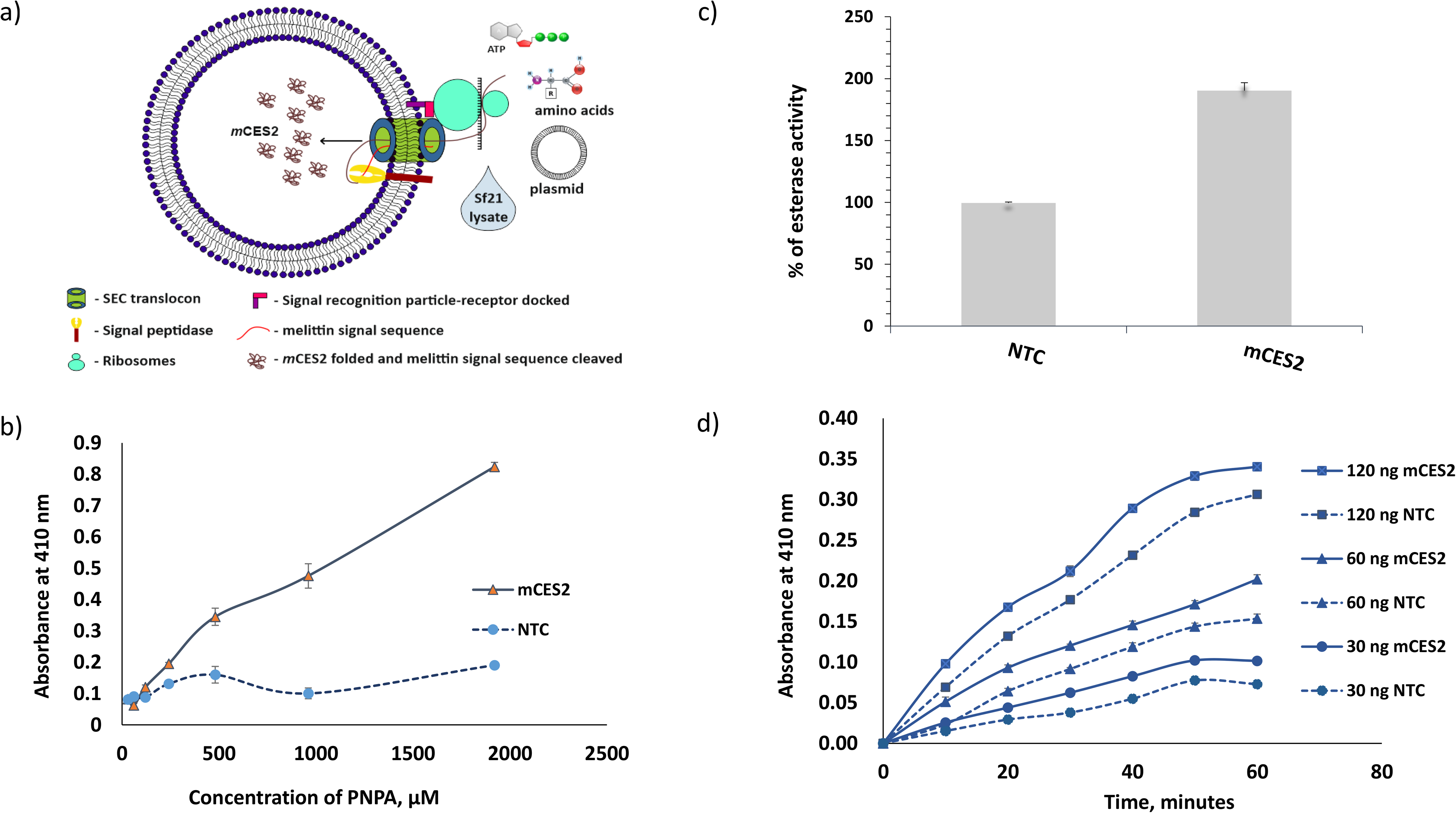
Esterase activity using PNPA. a) Schematic representation of cell-free synthesis of mCES2; b) Substrate concentration dependent esterase activity plot of *m*CES2, 37°C, 60 ng of mCES2, n = 3; c) Esterase activity of *m*CES2 using PNPA in the microsomes, 37°C, 60 min, 100 ng in 0.3 mM PNPA, n = 6; d) Time and enzyme concentration dependent plot of *m*CES2, 37°C with 0.3 mM PNPA, n = 3.

For calcium experiments, Fluo-5N AM was used due to the low binding affinity (90 μM) for Ca^2+^. The masked negative charge of the dye by acetoxymethyl ester moiety is unveiled with the aid of *m*CES2 inside the microsomes as shown in the schematic depiction Fig 3 a. For all experiments in the acetoxymethyl ester cleavage studies in Fig 3 b, c and d, no dye loading enhancers like pluronic F-127, probenecid or saponin were used due to their interference in fluorescence measurement caused by additive loading (25, 26). Both time dependent plot and substrate dependent plot showed a higher amount of dye loaded and cleaved for AM moiety. 5 μM of Fluo-5N AM was used for dye loading experiments measured by a microplate reader, 2 μM of Fluo-5N AM was enough for qualitative and quantitative measurements of calcium with confocal microscopy without any loading enhancers from *Sf*21 cell-free lysate. The *m*CES2 overexpression inside the microsomes significantly enhances the amount of dye loaded and reduces dye incubation time. As the *Sf*21 microsomes from the cell-free lysate undergo the translation reaction for 24 h at 30 °C, the esterase activity from the NTC microsomes is expected to be lower than the microsomes that are prepared for other purposes. Nevertheless, overexpression of *m*CES2 cuts down the concentration of Fluo-5N AM required for qualitative calcium measurements.

**Fig 3:**
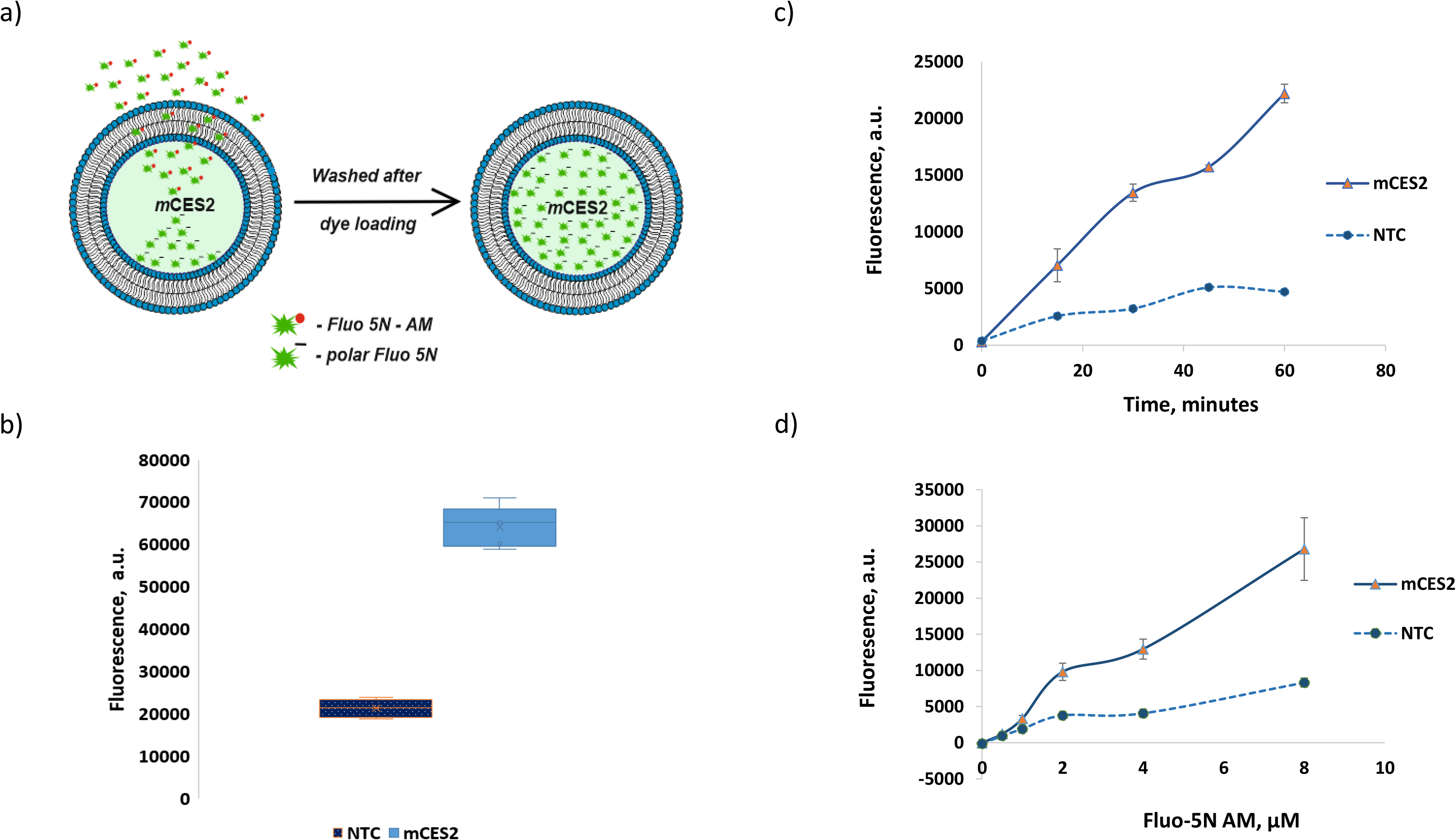
Calcium dye loading experiments. a) Illustration of calcium dye loading in *m*CES2 microsomes, b) Esterase activity performed with 200 ng of *m*CES2 each in 100 μl of 5 μM Fluo 5N-AM, 37°C, 60 min, n = 5; c) Time dependent esterase activity with 100 ng of *m*CES2 each in 100 μl of 5 μM Fluo 5N-AM, 37°C, n = 2; d) Fluo 5N-AM concentration dependent activity with 100 ng of *m*CES2 protein each in 100 μl reaction, 60 minutes, 37°C, n = 2.

### Establishment of TED based calcium imaging in microsomes

The K_d_ of Fluo-5N AM for *Sf*21 lysate microsomes was calculated to be 265 μM (Fig 4 a). The experiment was performed by first chelating resting calcium levels in the microsomes by 5 mM EGTA and then sequentially increasing the concentration of Ca^2+^ from 0 to 1 mM every step by 100 nM in the presence of 10 μM ionomycin. The K_d_ in phosphate buffer saline reported was 90 μM (27). The K_d_ *in vitro* or *in vivo* is usually higher than in the buffer which varies among different biological systems and experimental conditions. Sarcoplasmic reticulum vesicles for example from rabbit ventricular myoctes show a K_d_ of 400 μM (28) and mouse skeletal muscle sarcoplasmic reticulum have K_d_ of 133 μM (29). The K_d_ *in vitro* or *in vivo* is usually higher than in the buffer due to the varied expression of calcium binding proteins. In case of microsomes, it is due to the presence of calcium binding proteins derived from the sarcoplasmic reticulum like soluble calreticulin and single pass membrane protein calnexin that compete with Fluo-5N AM for luminal calcium (30).

**Fig 4:**
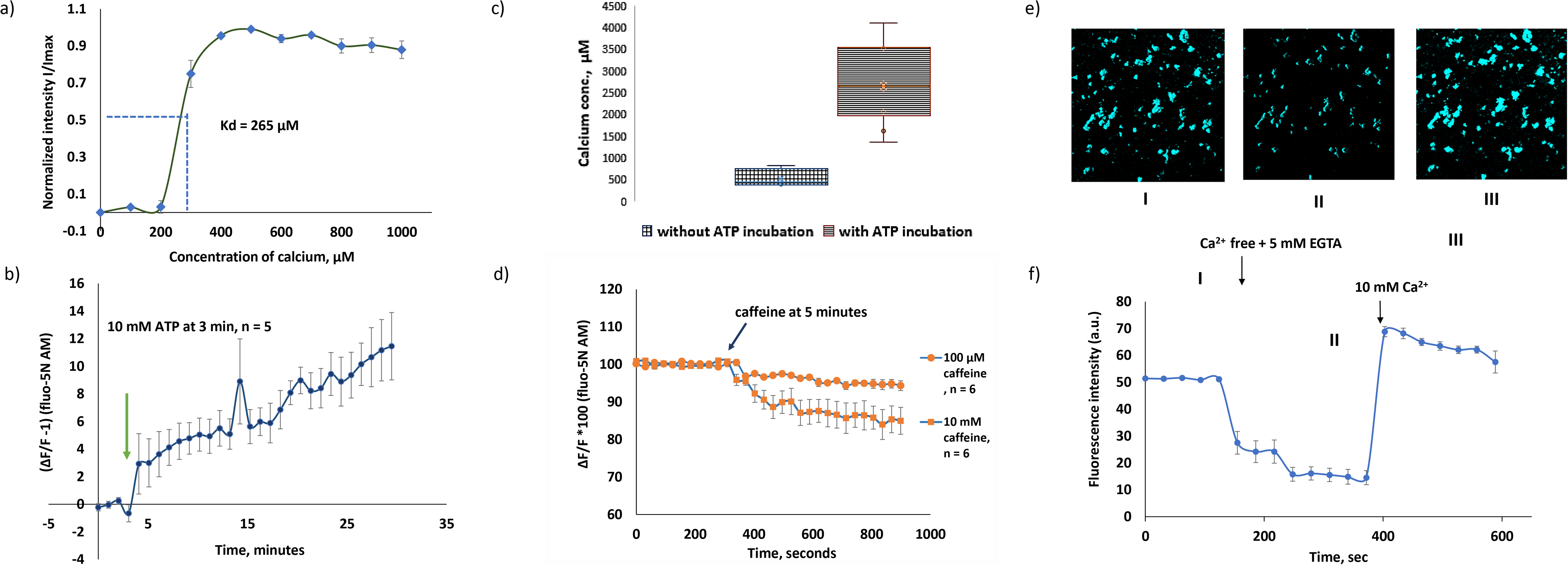
Direct calcium imaging using *m*CES2 microsomes. a) K_d_ evaluation of Fluo 5N-AM in the *Sf21* microsomes using 10 μM ionomycin plus varying concentration of calcium, n = 4, the data is represented as mean +/− S.E.M in the plot of I/I_max_ and Ca^2+^ concentration; b) SERCA pump activity in microsomes at room temperature with 10 mM ATP added at 3 min and measured for 30 minutes, n = 5. ATP induced calcium loading into the *m*CES2 microsomes was observed. The measurement was done using general calcium assay buffer plus the ATP regenerating system, 1 mM Mg^2+^ and 1 mM DTT; the data is represented as mean +/− S.E.M. The data before adding ATP is normalized to 0. c) Calcium levels comparison between ATP induced Ca^2+^ loaded and non-loaded microsomes at 37°C for 1 hour, n = 5 each, the data is represented as box plot with max and min; d) Caffeine induced calcium release in *m*CES2 microsomes via ryanodine receptor (RyR2) activation, n = 6 each for 10 mM and 100 μM caffeine. The data is represented as ΔF/F *100 of Fluo 5N-AM. Calcium was loaded in the *m*CES2 microsomes with ATP for 60 min, 37°C prior to caffeine experiments, e) Representative confocal images of I, II and III from different states of graph f); f) Representative graph of calcium concentration determination using Ca^2+^ free + 5 mM EGTA and 10 mM Ca^2+^ in the presence of 10 μM ionomycin.

Studying the ER resident proteins, using AM calcium dyes in the *m*CES2-expressive microsomes has several advantages over the endoplasmic reticulum in cell-based assays. As the low affinity AM dyes have to reach the endoplasmic reticulum through the cytosol where the carboxylesterases are abundant, a significant amount of fluorescent dye is cleaved and remain in the cytosol. Despite using low affinity Ca^2+^ indicators, which diminish the sensitivity to cytosolic Ca^2+^ levels, it will still contribute to the large noise relative to signal due to the high amount of cleaved dye present in the cytosol. Another advantage is that, it gives the flexibility to alter the calcium levels outside the microsomes, which could not be done likewise with cytosolic calcium levels. Largely, it aids in reducing the intrinsic calcium leakage due to the potentiation of Ca^2+^ between the luminal and outer face of the microsomes. It has already been proposed that by reducing the ER/SR calcium levels, one can reduce the calcium leakage of the SR in cells. In the case of ER derived microsomes, this is not necessary as we could increase the extra-microsomal Ca^2+^ to reduce the potentiation (31). For all calcium experiments, we used 200 μM calcium in the buffer in order to reduce the leakage from microsomes. Using 300 nM Ca^2+^ in the buffer caused extensive leakage relative to the 200 μM Ca^2+^ buffer as shown in supplementary data 2.

In order to study the native proteins in microsomes, extensively investigated proteins such as sarcoplasmic reticulum ATPase (SERCA) and ryanodine receptors were choice of interest. In *Sf*21 microsomes, SERCA activity was previously recorded with radiolabelled ^45^Ca^2+^ (32, 33). SERCA activity at room temperature was observed for about 30 min in the presence of 10 mM ATP with a slow increase in microsomal Ca^2+^ (Fig 4 b). Change in temperature alters the K_d_ with change in the sensitivity range and the fluorescence life time which in turn affects the bleaching rate of the Fluo-5N AM (34). As expected, we also observed that the K_d_ is lower and high bleaching was observed at 37°C compared to room temperature (data not shown). Hence, to avoid artifacts caused by temperature, we preferred to evaluate the increase in Ca^2+^ levels in microsomes at room temperature after treatment of microsomes at 37°C with ATP. SERCA activity cannot be monitored alone as Mg^2+^ ATPase also comes into play in the presence of ATP (35). Moreover, Mg^2+^ is necessary for SERCA activity and Fluo-5N AM binds to Mg^2+^ up to a certain extent with lower affinity relative to Ca^2+^ (Thermofisher Catalogue), it is to be observed that, SERCA activity measurement both indirectly by inorganic phosphate and directly by fluorescent indicators include Mg^2+^ ATPase activity as well. Microsomal preparations in general, include EDTA that can chelate both Ca^2+^ and Mg^2+^. Moreover, the cell-free lysate preparation procedure includes especially EGTA, a Ca^2+^ specific chelator for arresting the S7 nuclease treatment in the course of the process. On other hand, the calcium levels inside the microsomes could also be loaded up during the cell-free protein synthesis, as this reaction is performed with Mg^2+^ and 1 mM ATP at 30°C for 24 h. The typical [Ca^2+^] concentrations in the ER varies from few μM to mM (36). Calcium binding proteins like calreticulin, calnexin, GRP78 (BiP), GRP94, ERp72, protein disulfide isomerase, reticulocalbin, and ERC55 act as buffer causing the multiple order of variation in the [Ca^2+^] of the ER due to different individual binding affinities (36,37). The binding affinity for the Ca^2+^ buffering proteins are reported to be in the range of mM (38). In order to estimate the amount of calcium present in the microsomes, we have used used 10 μM ionomycin + 5 mM EGTA for F_max_, 10 μM ionomycin + 10 mM Ca^2+^ for F_min_ using the formula,

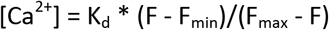

The representative graph for calcium concentration determination is shown in Fig 4 c and f. The sensitivity of the Fluo- 5N AM that can be visually observed is shown in Fig 4 e, representing I, II, III experimental conditions. With depletion of Ca^2+^ in microsomes, the decrease in fluorescence is noted in II and when the buffer is completely exchanged to 10 μM Ca^2+^, the increase in intensity is observed in III as recorded in video of Supplementary data 3. The estimated [Ca^2+^] in *Sf*21 microsomes that undergo CECF translation was estimated to be in the range of 100 to 1000 μM (Fig 3) which is coherent with previously reported data in other eukaryotic cells. Microsomes treated with 10 mM ATP, 1 mM Mg^2+^ with 200 μM Ca^2+^ for 1 h at 37 °C, showed a 4-5 times higher range of luminal Ca^2+^. The in-vitro K_d_ which we estimated is 265 μM, and it is approximately 3X higher compared to the K_d_ in the buffer solution. The sensitivity range or dynamic range of Fluo-5N vary from 10^−7^ to 10^2^ M of Ca^2+^ *in vitro* for the sarcoplasmic reticulum (29). Microsomes treated for SERCA activity showed a median value of 2500 μM which is in accordance with the expected sensitivity range (Fig 4 c).

Ryanodine receptors, RYR1 and RyR2 perform a plethora of functions in mammalian physiology ranging from skeletal muscle and cardiac muscle contraction, cognitive functions such as learning and memory. Ryanodine channels that deplete the calcium stores in response to ryanodine are ER resident leak channels, RyRs respond to ryanodine at low concentration and caffeine. Ryanodine receptors have been extensively investigated in eukaryotic cells (39, 40, 41, 42). Fig 4 d represents the calcium response of 100 μM and 10 mM caffeine. 10 mM caffeine induced a higher calcium efflux relative to 100 μM caffeine.

### Analysis of cell-free synthesized *h*TRPV1 channel

In pursuance of calcium imaging with cell-free synthesized channels, we have chosen the human Transient Receptor Potential Channel, Vallinoid Receptor member 1 (*h*TRPV1) as a model protein. *h*TRPV1 expressed in cells in the plasma membrane has been studied for calcium entry into cytoplasm using fura 2-AM (43, 44).

*h*TRPV1 is largely present in the plasma membrane with the large N terminal and the C terminal stretches towards the cytosol in cells. As expected, in cell-free synthesis, the orientation of the protein incorporated in the microsome is opposite to that of the native plasma membrane orientation. Despite most of the activators and inhibitors for the TRP channel family are organic membrane permeable compounds, attention should be paid while working with the non-polar activators or inhibitors. All experiments depicted in Fig 5 are performed after calcium loading into the microsomes for 1 h in order to ensure higher [Ca^2+^] inside the microsomes. With the above-mentioned experimental conditions, we have observed that the activation of the *h*TRPV1 caused Ca^2+^ release from *h*TRPV1-*m*CES2 microsomes. Saturation of capsaicin induced Ca^2+^ release was observed even with 200 nM of CAP in the *h*TRPV1-*m*CES2 microsomes. The Ca^2+^ release by 200 nM CAP was abolished completely 20 μM capsazepine (CPZ) (Fig 5 c). The concentration for activation and inhibition of *h*TRPV1 in our experiments are coherent with the molar range as the cell-based expression (45). Presence of Phosphatidylinositol 4,5-bisphosphate, (4,5)PIP_2_ for activation of TRP channels has been debated by several researchers (46, 47, 48). In our work, we have shown that without external (4,5)PIP_2_, activation of TRPV1 was feasible. On the other hand, the phenomenon we observe can be supported by the presence of (4,5)PIP_2_ in the endoplasmic reticulum and in the cytosol (48, 49, 50).

**Fig 5:**
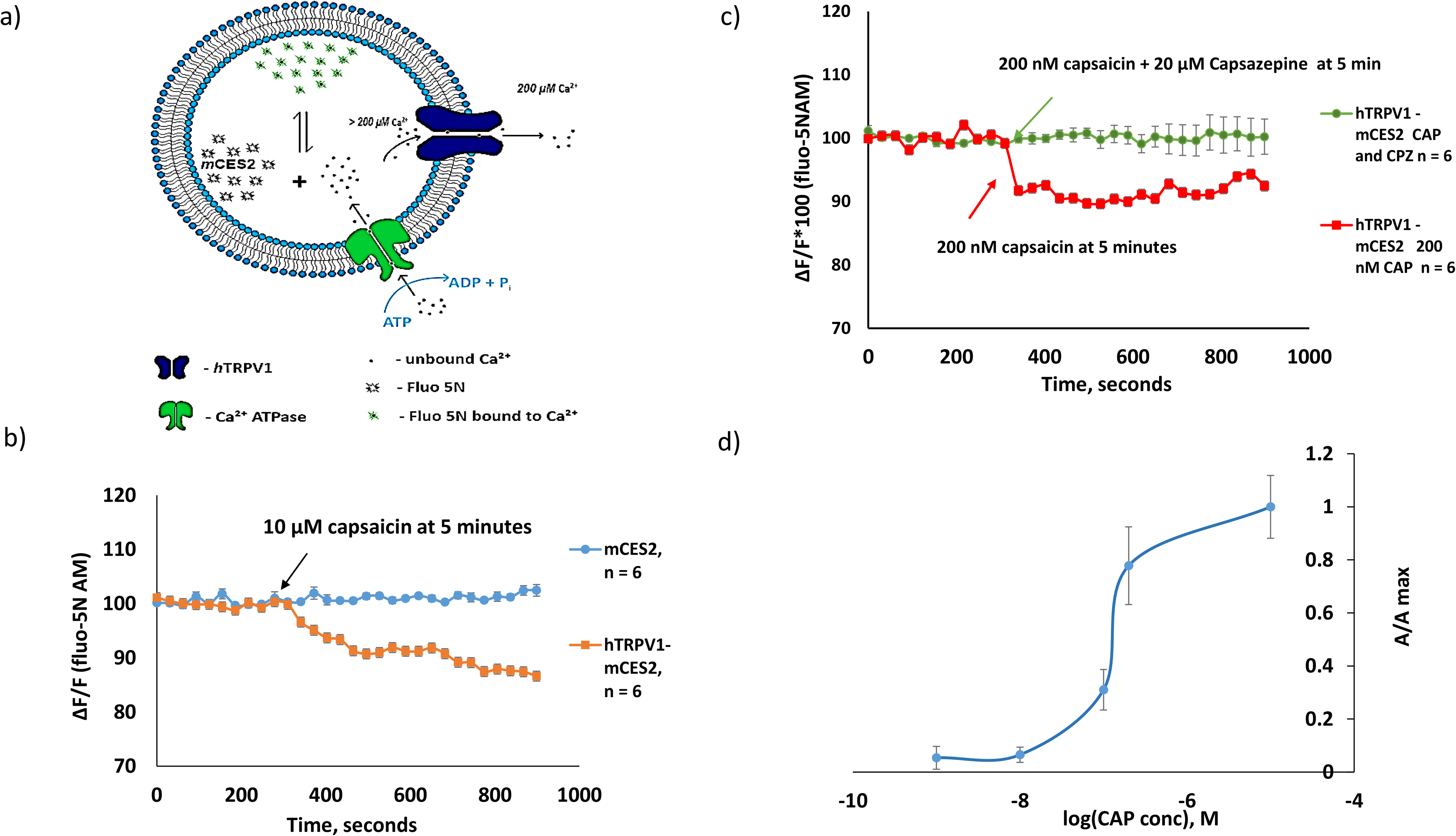
Functional assessment of cell-free synthesized *h*TRPV1. a) Illustration of calcium dynamics in microsomes expressed with *h*TRPV1 and *m*CES2; b) Comparison plot of *h*TRPV1-*m*CES2 and *m*CES2 microsomes for capsaicin (CAP) induced Ca^2+^ release from microsomes, n = 6 each. 10 μM capsaicin was used at 5 min and the release was monitored further for 10 minutes. No Ca^2+^ release was observed in the *m*CES2 microsomes; c) Capsazepine mediated inhibition of calcium release caused by capsaicin in *h*TRPV1-*m*CES2 microsomes, 200 nM CAP and 20 μM CPZ were used, n = 6 each. All samples in c) and d) were calcium loaded for 60 min, 37°C using ATP before start of capsaicin and capsazepine studies. All data in b) and c) is represented as ΔF/F*100 of baseline before adding the stimulant; d) Dose dependent response of capsaicin in *h*TRPV1-*m*CES2 microsomes, n = 4 to 7 per each data point.

## CONCLUSION AND OUTLOOKS

Apart from the TED method, which we utilize in our work, also other fluorescent methods can be employed. For instance, using membrane impermeable Ca^2+^ dyes to measure the extra-microsomal solution, which works conversely to the TED -based method (51). But with this method, only absolute luminal calcium levels cannot be monitored. Only the fluctuation caused by calcium influx/efflux can be measured. Radioactive calcium measurements have widely been used by researchers for studying microsomes (13, 14, 15, 16). Major drawback when studying microsomes with ^45^Ca^2+^ is that, first the microsomes should be loaded with ^45^Ca^2+^ using SERCA for efflux studies. Without ^45^Ca^2+^, only influx studies can be performed. In the above-mentioned methods, as extra microsomal calcium is monitored and the microsomes are not immobilised on the microplate/cuvette, live monitoring of Ca^2+^ is impracticable while exchanging activator and inhibitor solutions. Moreover, endpoint-based methods increase the material costs while studying the kinetics. As chemical calcium dyes are toxic upon prolonged usage in live cells or animal models, genetically encoded calcium indicators for ER, mitochondria and other organelles are new emerging areas of research since the development of genetically engineered D1ER from Roger.Y. Tsien group (52). Though, genetically encoded calcium dyes could be another alternative other than the TED based system, the practicability of D1ER expressed cell line for CFPS preparation should be tested. Membrane bound ER proteins are well retainable in microsomes. However, loss of lumen soluble proteins is expected on due course of cell-free lysate preparation.

Apart from calcium imaging in a CFPS platform using a chemical fluorescent dyes, there are other biological applications for carboxylesterases. Carboxylesterase have known to impart pesticide resistance in insects (53, 54). Carboxylesterases play an important role in xenobiotic metabolism of environmental toxins and drugs. Moreover, several human drugs are consumed as pro-drugs which are further processed in liver by carboxylesterases into active drug formulation (55,56). In this work, we have shown that the functionally active carboxylesterases could be synthesized using CFPS platform. As the synthesized carboxylesterase is present in microsomal lumen, active transport of prodrugs across the microsomal membrane can also be monitored. Hence, in order to cope up with the sustenance needs of above-mentioned areas of carboxylesterase research, cell-free synthesized carboxylesterases could be a conducive platform thereby surpassing the conventional disadvantages of cell-based protein synthesis.

## MATERIALS AND METHODS

### Continuous Exchange Cell-Free (CECF) translation

Eukaryotic CECF translation of proteins were performed using *Sf*21 lysates in the special dialysis chamber containing two compartments separated by 10kDa cut off dialysis membrane in between reaction mixture and the feeding mixture. A 50 μL standard reaction mixture of a *Sf*21 cell-free synthesis reaction in the reaction chamber was composed of 40% lysate, 30 mM HEPES-KOH, 2.5 mM Mg(OAc)_2_, 75 mM KOAc, 0.25 mM spermidine, 100 μM of each canonical amino acid, nucleoside triphosphates (1.75 mM ATP, 0.30 mM CTP, 0.30 mM GTP, and 0.30 mM UTP), 120 ng/μl plasmid DNA, and 1 U/ml T7 RNA-polymerase, 20 μM of PolyG, 30 μM of the caspase inhibitor -Z-VAD-FMK (benzyloxycarbonyl-Val-Ala-Asp(OMe)-fluoromethylketone), and 0.02% of sodium azide. The feeding mixture of 1 ml contained all the above components except plasmid, PolyG, T7 RNA polymerase and *Sf*21 lysate. ^14^C-labeled leucine (100 dpm/pmol) for the detection of *de novo* synthesized proteins. For functional analysis, the proteins were synthesized in the absence of ^14^C-leucine. Protein translation reactions based on *Sf*21 lysates were incubated for 24 hrs at 30 °C, 600 rpm using a thermomixer (Eppendorf, Hamburg, Germany). If required translation mixture (TM) of cell-free reactions were further fractionated for analysis. The fractionation was realized by centrifugation at 16,000 g for 10 min at 4°C in order to separate the ER-derived vesicular fraction (VF) of the cell lysate from the supernatant (SN). The microsomal fraction was suspended in PBS buffer without calcium and magnesium ions for further analysis such as quantification of protein yields. For storage, the total translation mix is snap frozen in liquid nitrogen and stored at −80°C.

### Quantification of Cell-Free Synthesized Protein Yields

Based on the incorporation of ^14^C-leucine in cell-free synthesized proteins, the respective protein yield can be estimated by scintillation measurement. Therefore, 5 μL aliquots of each translation mixture were mixed with 3 mL of a 10% (v/v) TCA-2% (v/v) casein hydrolysate solution in a glass tube and incubated at 80 °C for 15 min. Afterwards, samples were chilled on ice for 30 min and retained on the surface of glass fiber filter papers using a vacuum filtration system. Filter papers were washed twice with 5% TCA and then vacuum dried with acetone. Dried filters were placed into a scintillation vial, 3 mL of scintillation cocktail was added and vials were agitated on an orbital shaker for at least 1 hr. The scintillation signal was determined using the LS6500 Multi-Purpose scintillation counter. The protein yields were identified based on the obtained scintillation counts and protein specific parameters including molecular mass and amount of leucine. Error bars calculated for the protein yield show the individual standard deviation.

### SDS-PAGE and Autoradiography

The molecular size of radiolabeled, cell-free synthesized protein was analyzed using SDS-PAGE followed by autoradiography. First, 5 μL of the respective fraction of a cell-free synthesis reaction including the radiolabeled target protein was subjected to ice-cold acetone. Precipitated protein was separated by centrifugation (16,000 × g, 4 °C, 10 min) and the protein pellet was dried for at least 30 min at 45°C. The dried protein pellet was dissolved in LDS sample buffer with 50 mM DTT and loaded on a pre-casted NuPAGE 10% Bis-Tris gel. The gel was run at 185 V for 35 min according to the manufacturer’s protocol. Subsequently, gels were dried at 70 °C, placed on a phosphor screen and radioactively labeled proteins were visualized using a Typhoon Trio + variable mode.

### Esterase activity using para-nitrophenol

To determine the esterase activity, m-CES2 was translated using *Sf*21 CECF reaction. 50 μl of total translation mix was first centrifuged at 16,000 × g for 10 minutes at 4°C. The microsomal pellet was again washed with Phosphate Buffer Saline (PBS) with no Ca^2+^ and Mg^2+^ to remove the native cytosolic carboxylesterases outside the microsomes from the lysate via further centrifugation. The pellet was dissolved in esterase assay buffer containing 20 mM Tris–HCl (pH8.0), 150 mM NaCl, and 0.01% Triton X-100. Fresh solutions of 4-para-nitrophenylacetate (PNPA) were used as the substrate. 250 μl of PNPA substrate solution of 150 μM concentration was used to initiate the reaction and the mixture was incubated at 37 °C for 1 hour. The para-nitro phenol formed after esterase activity was measured using a Mithras Plate reader at 410 nm. No protein was added for blank reactions and the esterase activity was evaluated as percentage of NTC samples.

### Dye loading assays with Fluo-5N AM

50 μl of translation mix was first centrifuged 16,000 g for 10 minutes at 4°C to obtain the microsomal pellet. The microsomes were resuspended in ATP based calcium imaging buffer to initiate the SERCA activity. The calcium imaging buffer was composed of 75 mM KCl, 20 mM HEPES-KOH, 5 mM NaN_3_ and 200 μM CaCl_2_ with pH 7.4. To enhance SERCA activity, 10 mM adenosine 5’ triphosphate (ATP), 1 mM MgCl_2_, 0.5 mM dithiothreitol (DTT), 5 mM phosphocreatine (PCr), and 20 U/mL creatine phosphokinase (CPK). SERCA activity induced Ca^2+^ loading was performed at 37°C for 30 minutes and then stopped by centrifuging and removing the supernatant. The pellet is suspended in 100 μl of Fluo-5N AM dye of concentration 2 μM and then incubated at 37°C at 500 rpm. The reaction is stopped by centrifugation and removal of the dye. Microsomes are further washed with 100 μl of PBS to purge the uncleaved Fluo-5N AM from the microsomes at 37°C for 20 minutes and centrifuged to obtain the microsomal pellet containing only the cleaved dye. Then, the pellet washed and fluorescence removed subsequently was measured in plate reader with E_x_ 488 and E_m_ 515 nm. The blank sample measurements were subtracted and the data is analyzed.

### Calcium imaging using confocal laser microscopy

For all calcium measurements, the microsomes were seeded for attachment on the coverslip coated with poly D lysine Hydrobromide. The coverslips were autoclaved and coated overnight with poly-D-lysine Hydrobromide (0.1 mg/ml), dried and stored at room temperature. The microsomes were seeded for 1 h, 37°C and the coverslips were washed with the calcium imaging buffer for removal of unsettled microsomes. A flow chamber fitting the coverslip at the bottom with the aid of vacuum sealing agent was used for all measurements. Argon laser with Alexa 488 at 3% intensity, maximum gain and 5-6 airy units were used for all measurements using the LSM Meta 510 software (Carl Zeiss Microscopy, GmbH) time series function with a frequency of 30 s and a 40X oil immersion objective with numerical aperture 1.3 was used. The data is represented as delta F/F, where delta F represents the difference in fluorescence of microsomes and background. Different regions of interest were selected from each individual experiment from the frame of 512 μM X 512 μM frame for observing whether identical increase or decrease in intensity is present. The slope of bleaching was drift corrected using peak and baseline correction protocol using the OriginPro 2015 software. For comparison of individual experiments with activators and inhibitors, the baseline fluorescence intensity was normalized to 100 a.u. and then increase or decrease in intensity was analyzed. Statistical analysis and plotting graphs were performed either by OriginPro and Microsoft Excel software.

**Table 1:** Esterase activity of mCES2 (mouse Carboxyl ESterase 2) and NTC (Non-Template Control) samples using pNPA method. Data table of Fig 2 d representing time dependent and dose-dependent curve shown above as absorbance at 410 nm.

|  | *m*CES2 samples |  |  |  |  |  |
| --- | --- | --- | --- | --- | --- | --- |
|  | 30 ng |  | 60 ng |  | 120 ng |  |
| Time (sec) | average | S.E.M | average | S.E.M | average | S.E.M |
| 0 | 0.000000 | 0.000000 | 0.000000 | 0.000000 | 0.000000 | 0.000000 |
| 10 | 0.025467 | 0.000720 | 0.051200 | 0.002265 | 0.097800 | 0.003859 |
| 20 | 0.043867 | 0.001785 | 0.093000 | 0.002868 | 0.167533 | 0.004907 |
| 30 | 0.062200 | 0.003559 | 0.120333 | 0.001785 | 0.211867 | 0.006706 |
| 40 | 0.082533 | 0.000720 | 0.145667 | 0.013411 | 0.289200 | 0.003266 |
| 50 | 0.102000 | 0.001633 | 0.171267 | 0.014350 | 0.329000 | 0.005354 |
| 60 | 0.101467 | 0.000544 | 0.201867 | 0.014631 | 0.340467 | 0.001361 |
|  | NTC samples |  |  |  |  |  |
|  | 30 ng |  | 60 ng |  | 120 ng |  |
| Time (sec) | average | S.E.M | average | S.E.M | average | S.E.M |
| 0 | 0.000000 | 0.000000 | 0.000000 | 0.000000 | 0.000000 | 0.000000 |
| 10 | 0.015133 | 0.002880 | 0.023467 | 0.005761 | 0.069133 | 0.001515 |
| 20 | 0.029200 | 0.001247 | 0.064200 | 0.003399 | 0.131867 | 0.004481 |
| 30 | 0.037867 | 0.003954 | 0.091533 | 0.001785 | 0.176867 | 0.003839 |
| 40 | 0.054867 | 0.004119 | 0.118867 | 0.004907 | 0.231533 | 0.003067 |
| 50 | 0.077333 | 0.003839 | 0.143667 | 0.004380 | 0.284333 | 0.002373 |
| 60 | 0.072800 | 0.000817 | 0.153467 | 0.005664 | 0.306467 | 0.003345 |

## ACKNOWLEDGMENTS

This work was supported by German Ministry of Education and Research, BMBF (BMBF, No. 031B0078A). We would like to appreciate Dr. Rita Sachse for her preliminary attempts in this topic during her Ph.D thesis in our lab. We would like to acknowledge Ms. Dana Wenzel (Fraunhofer IZI, Potsdam-Golm, Germany) for technical support, Dr. Lena Thoring and Dr. Marlitt Stech (Fraunhofer IZI, Potsdam-Golm, Germany) for providing their insights and discussion in the cell-free synthesis platform in due course of this research work. We would also like to thank Dr. Michael Kirschbaum for helping us with the imaging platforms.

## AUTHOR CONTIBUTIONS

PD and SK have framed the basic idea for this work. D A.W was responsible for production and quality control of cell-free lysates used in this manuscript. AZ, DB and PD have performed experiments. PD, SD and SK have contributed in preparing the manuscript.

---

**Supplementary data 2:**
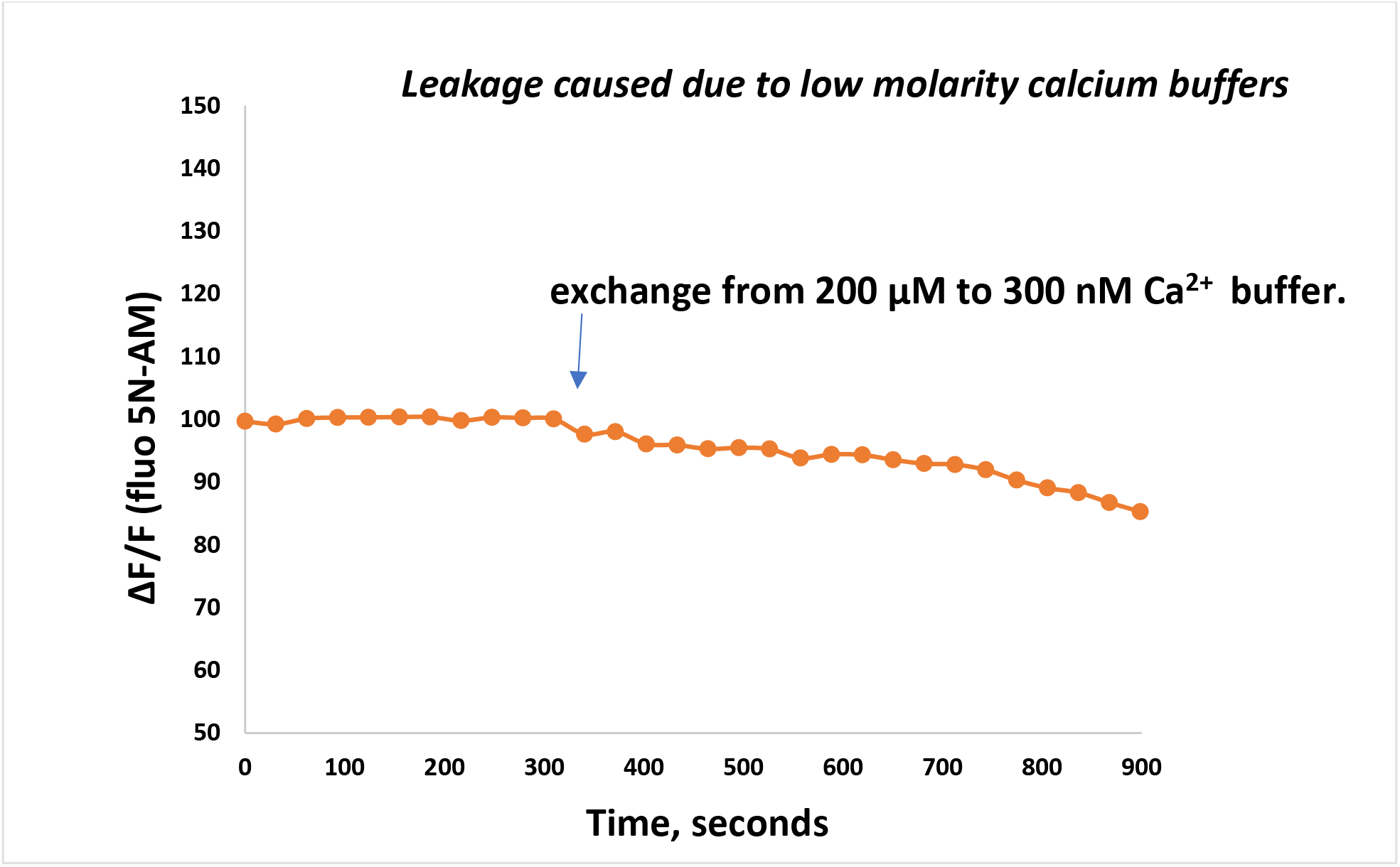
Gradual leakage caused due to change in the buffer from 200 μM to 300 nM.

